# Spatial proteomics of *Onchocerca volvulus* with pleomorphic neoplasms shows local and systemic dysregulation of protein expression

**DOI:** 10.1101/2024.10.15.618383

**Authors:** Lucia S. Di Maggio, Kerstin Fischer, Bruce A. Rosa, Devyn Yates, Byoung-Kyu Cho, Jessica Lukowski, Antonia Zamacona Calderon, Minsoo Son, Young Ah Goo, Nicholas O. Opoku, Gary J. Weil, Makedonka Mitreva, Peter U. Fischer

**Affiliations:** Infectious Diseases Division, Department of Medicine, Washington University School of Medicine, St. Louis, MO 63110, USA; Mass Spectrometry Technology Access Center at McDonnell Genome Institute, Washington University School of Medicine, St. Louis MO 63110; Fred Newton Binka School of Public Health, University of Health and Allied Sciences, Ho, Ghana; McDonnell Genome Institute, Washington University School of Medicine, St. Louis, MO 63108, USA; Department of Genetics, Washington University School of Medicine, St. Louis, MO 63110, USA

## Abstract

*Onchocerca volvulus* is the agent of onchocerciasis (river blindness) and targeted by WHO for elimination though mass drug administration with ivermectin. A small percentage of adult worms develop pleomorphic neoplasms (PN) that are positively associated with the frequency of ivermectin treatment. Worms with PN have a lower life expectancy and a better understanding about the proteins expressed in PN, and how PN affect protein expression in different tissues could help to elucidate the mechanisms of macrofilaricidal activity of ivermectin. Within a clinical trial of drug combinations that included ivermectin, we detected 24 (5.6%) *O. volvulus* females with PN by histology of paraffin embedded nodules. To assess the protein inventory of the neoplasms and to identify proteins that may be associated with tumor development, we used laser capture microdissection and highly sensitive mass spectrometry analysis. Neoplasm tissue from three female worms was analyzed, and compared to normal tissues from the body wall, uterus and intestine from the same worms, and to tissues from three females without PN. The healthy females showed all intact embryogenesis. In PN worms, 151 proteins were detected in the body wall, 215 proteins in the intestine, 47 proteins in the uterus and 1,577 proteins in the neoplasms. Only the uterus of one PN female with some stretched intrauterine microfilariae had an elevated number of proteins (601) detectable, while in the uteri of the healthy females 1,710 proteins were detected. Even in tissues that were not directly affected by PN (intestine, body wall), fewer proteins were detected compared to the corresponding tissue of the healthy controls. Immunolocalization of the calcium binding protein OvDig-1 (OVOC8391) confirmed the detection in PN by mass spectrometry. In conclusion we identified proteins that are potentially linked to the development of PN, and systemic dysregulation of protein expression may contribute to worm mortality.

**Author summary:** *Onchocerca volvulus*, the causative agent of onchocerciasis (river blindness), is targeted for elimination by WHO. The primary strategy involves mass administration of ivermectin. A small proportion of adult female worms develop pleomorphic neoplasms (PN). Here, we used laser capture microdissection and highly sensitive mass spectrometry analysis to determine the protein inventory of PN to identify proteins that may be associated with tumor development. Neoplasm tissue from female worms was analyzed, and compared to normal tissue from the body wall, uterus and intestine from the same worms, and to tissues from females without PN. When compared, PN and healthy control (HC) worms display a different set of proteins, the PN tissue being the one with the highest number of proteins (1,390). From these, 594 were not present in any HC worm tissue. Despite the large number of proteins identified in PN tissue, their low abundance suggests also in PN dysregulation of protein expression. Immunolocalization of a calcium binding protein detected in PN confirmed the mass spectrometry results. In conclusion, we have developed a system to analyze the proteome of *O. volvulus* from nodule sections and identified proteins that are potentially linked to the development of PN and may contribute to worm mortality.

## Introduction

*Onchocerca volvulus* is a filarial nematode parasite and the agent of onchocerciasis, also known as river blindness. Onchocerciasis is a neglected tropical disease that is targeted for global elimination. Recent data from the World Health Organization (WHO) estimated that over 220 million people live in areas at risk of onchocerciasis transmission, almost exclusively in sub-Saharan Africa (1). Adult worms reside in subcutaneous nodules (onchocercomas) and have a reproductive life span of up to 15 years. The female worms release millions of microfilariae that migrate through the skin and sometimes invade the eye and may cause eye disease. Male worms grow up to 8 cm and migrate from one nodule to another, while females can grow up to 60 cm and are permanently coiled up within the nodules. Nodules vary in size from less than 1 cm to up to 5 cm in diameter and can contain up to 12 females, although about 30% of the nodules contain only a single female (2).

Currently there is no drug available that efficiently kills all adult worms that can be used for mass drug administration (MDA) (3). The main strategy to eliminate onchocerciasis is annual or semiannual MDA with ivermectin alone or ivermectin plus albendazole in areas co-endemic for lymphatic filariasis. Ivermectin efficiently kills microfilariae and after repeated doses it also has a limited effects on adult worms. A small percentage of older, adult female *O. volvulus* are known to develop pleomorphic neoplasms (PN), and it appears that this percentage increases after several doses of ivermectin and can be as high as 10% (4, 5). The induction of PN has been considered as one mechanism how ivermectin contributes to the death of worms as the filarial origin of the neoplasms and the pleomorphism of the tumor cells have been addressed in previous studies (4, 6). Morphological studies have shown that female worms with PN have abnormal embryogenesis, are often sterile and do not produce viable microfilariae. This sterility is independent of the location of the PN, and it is independent whether they are confined to the pseudocoelomic cavity or are within the reproductive system. PN tissue can be detected in distal and proximal parts of a female, indicating that neoplasms can reach many cm in length. Immunohistochemical studies have shown that PN consist of several different cell types that can be labeled with various antibodies raised against worm proteins (4). Although it assumed that the ovaries are the origin of the PN, the protein inventory of PN is unknown.

Technical advances in proteome analysis by liquid chromatography mass spectrometry (LC/MS) and increased genome information of *O. volvulus* together with improved laser capture microdissection (LCM) have made visual proteomics of *O. volvulus* possible. Visual proteomics combines morphology and digital image analysis with LCM and ultra-high-sensitivity LC/MS (7). If a standard amount of tissue is studied, the method can provide semi-quantitation of proteins. Several studies describe the proteome of whole adult female *O. volvulus* worms, and between 2,100 and 3,900 expressed proteins have been identified in this lifecycle stage (8–10). However, these studies used adult female worms of different ages that contain various reproductive stages, and no proteomic information is available at the tissue or organ level of individual worms.

The objective of the current study was to establish a protocol for efficient visual proteomic analysis of paraffin-embedded *O. volvulus* nodules. We compare the proteomic inventory of PN tissue to the inventory of the body wall (hypodermis, lateral chords, muscles), uterus and intestine from three worms with PN and three worms without neoplasms with intact embryogenesis. We were able to show that ethanol fixed, paraffin embedded nodules are a rich source for proteomic analysis of adults *O. volvulus*; more than 2,200 proteins were identified in the analyzed tissue types. We identified a specific proteomic inventory of proteins in PN that could be associated with the development of tumors in *O. volvulus*.

## Materials and methods

### Ethics statement

The protocol for the clinical trial was reviewed and approved by ethical review committees at the University of Health and Allied Sciences (UHAS) in Ho, Ghana, the Ghana Health Service, The Ghana Food and Drug Authority, Case-Western Reserve University (Cleveland, OH, USA) and Washington University School of Medicine (St. Louis, MO, USA) (IRB ID: 201910085) (11). The use of de-identified nodule sections for further analysis is not considered human subjects research according to the institutional review of Washington University School of Medicine.

### Nodule samples preparation for laser capture microdissection

*O. volvulus* nodules were obtained from a clinical trial that compared the tolerability and efficacy of IDA (ivermectin, diethylcarbamazine, albendazole) versus a comparator treatment (ivermectin plus albendazole) in persons with onchocerciasis in the Volta region, Ghana (11). For LCM, six nodules from female worms (three with PN, three healthy controls (HC)) were selected with biological triplicate samples of each condition (Table 1). PN female worms represented 5.6% (24/428) of all female adult worms found in participants from the Ghana trial (Table 2, see ref 10 table 4). For each parasite, four body regions were selected: body wall (including hypodermis, lateral chord and muscles), gut/intestine, uterus and neoplasm tissue or embryos (morulae, coiled and stretched microfilariae) (Figure 1).

**Fig 1.**
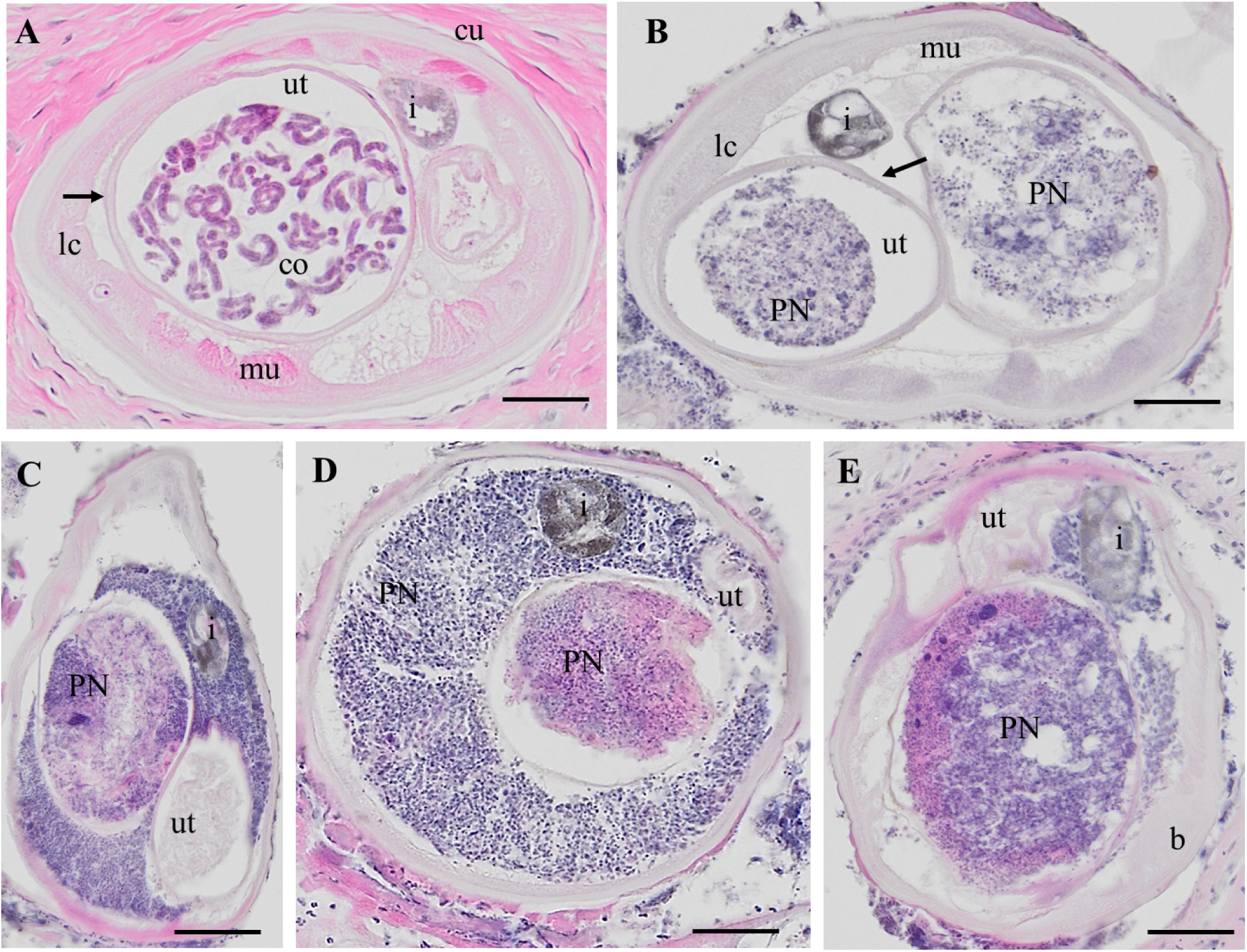
H&E stained cross-sections of *O. volvulus* females. A: HC female with coiled embryos in the uterus. B: PN worm with neoplasm in both uterus branches. C: PN female worm with neoplasm in one uterus branch and in the pseudocoelomic cavity. The second uterus branch is filled with degenerated embryos and was not used for LCM. D: PN female with neoplasm in one uterus branch and the pseudocoelomic cavity. The second uterus branch is empty. E: PN cells in one uterus branch and in the pseudocoelomic cavity. HC: healthy control, PN: pleomorphic neoplasm, ut,, b= body wall, co= coiled embryos i = intestine, lc = lateral chord, mu= muscle, ut =uterus, scale bar= 50 μm

**Table 1.**
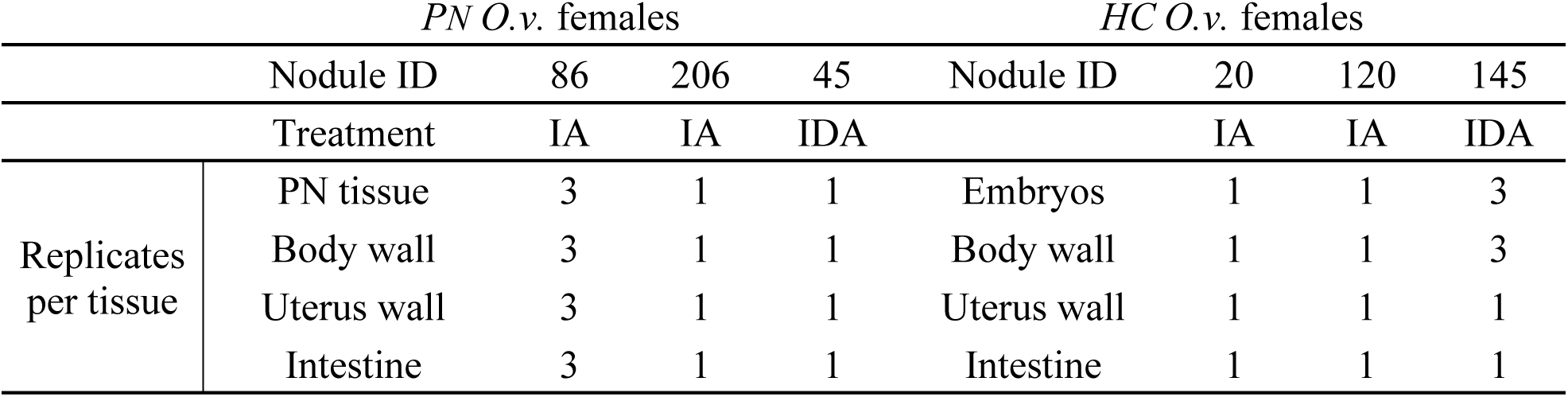
Sample information for the paraffin-embedded *O. volvulus* nodules. . Three different PN and three healthy control (HC) worms were selected from the IDA trial, Ghana. Four body regions were selected for laser capture microdissection: body wall (including hypodermis, lateral chord and muscles), intestine, uterus wall and PN tissue or embryos (morulae, coiled and stretched mf). I: ivermectin, D: diethylcarbamazine, A: albendazole and PN: pleomorphic neoplasm.

**Table 2.** Summary of nodule data by treatment group. The table shows the total number of nodules evaluated 18 months after treatment, summarizing the total number of alive females and number of alive PN females in each treatment arm. I: ivermectin, D: diethylcarbamazine and A: albendazole and PN: pleomorphic neoplasm.

|  | IA (%) | IDA (%) | IDA3 (%) | Total (%) |
| --- | --- | --- | --- | --- |
| No. nodules (evaluated) | 116 | 149 | 167 | 432 |
| No. living female worms | 127 | 142 | 159 | 428 |
| No. PN females (%) | 8 (6.3) | 4 (2.8) | 12 (7.5) | **24 (5.6)** |

From the collection of available *O. volvulus* nodules, the ones with only one living female were selected. For the LCM, hand-color coded images from either hematoxylin and eosin (H&E) or *O. volvulus* aspartic protease (Ov-APR)-stained sections were used to avoid potential contamination with sperm or dead or degenerate embryos, which can also be found in the uteri (S1 Fig.) (12). At least ten worm sections needed to be available for LCM for each female. Three slides with three 10 μm sections were prepared on PEN (polyethylene naphthalate) slides for each nodule. Stained parallel sections of each nodule were analyzed by microcopy and guided the tissue identification of the unstained LCM material. Tissue was collected from three nodule sections to assure sufficient material using the LCM System LMD7000 (Leica Microsystems GmbH, Wetzlar, Germany). Tissue was compiled into the cap of a 0.5 mL PCR tube (Axygen, Union City, CA, USA) containing 30 μL of 8M urea buffer. Collection tubes were spun down and ready for subsequent preparation for mass spectrometry analysis.

### Sample preparation for LC-MS/MS

The collected LCM cells were lysed in 8M urea-containing buffer. Samples were then reduced with 4 mM dithiothreitol (DTT) at 50°C for 30 minutes, followed by alkylation of cysteine residues with 18 mM iodoacetamide for 30 minutes in the dark. Proteins were digested with 2 µg of trypsin overnight at 37°C. The resulting peptides were desalted using solid-phase extraction with a C18 spin column and eluted with 0.1% trifluoroacetic acid (TFA) in 50% acetonitrile (ACN). Peptides from biological replicates (three PN and three HC worms) were reconstituted in 0.1% formic acid (FA) in water and analyzed by LC-MS/MS using a Vanquish Neo UHPLC system coupled to an Orbitrap Eclipse Tribrid Mass Spectrometer with a FAIMS Pro Duo interface (Thermo Fisher Scientific, San Jose, CA). Samples were loaded onto a Neo trap cartridge coupled with an analytical column (75 µm ID x 50 cm PepMapTM Neo C18, 2 µm) and separated using a linear gradient of solvent A (0.1% FA in water) and solvent B (0.1% FA in ACN) over 120 minutes. For MS acquisition, FAIMS switched between CVs of −35 V and −65 V with cycle times of 1.5 s per CV. MS1 spectra were acquired at 120,000 resolution, with a scan range from 375 to 1500 m/z, AGC target set at 300%, and maximum injection time set at Auto mode. Precursors were filtered using monoisotopic peak determination set to peptide, charge state 2 to 7, dynamic exclusion of 60 s with ±10 ppm tolerance. For the MS2 analysis, the isolated ions were fragmented by assisted higher-energy collisional dissociation (HCD) at 30% and acquired in an ion trap. The AGC and maximum IT were Standard and Dynamic modes, respectively. Data were searched using Mascot (v.2.8.3 Matrix Science) against two sets of combined databases. The first set is including *Homo sapiens*, cRAP, and *O. volvulus* databases, and the second set is *Wolbachia* and virus databases only. Trypsin was selected as the enzyme, and the maximum number of missed cleavages was set to 3. The precursor mass tolerance was set to 10 ppm, and the fragment mass tolerance was set to 0.6 Da for the MS2 spectra. Carbamidomethylated cysteine was set as a static modification, and dynamic modifications were set as oxidized methionine, deamidated asparagine/glutamine, and protein N-term acetylation. The search results were validated with 1% FDR of protein threshold and 90% of peptide threshold using Scaffold v5.3.0 (Proteome Software Portland, OR, USA). Proteins containing indistinguishable peptides based on MS/MS analysis alone were grouped to satisfy the principles of parsimony. Two independent Scaffold files were generated for *O. volvulus* and Wolbachia data sets. Spectra counts and peptide count for each peptide sequence are provided in Excel spreadsheets (Microsoft, Redmond, WA, USA) (Tables S1 and S2). The mass spectrometry proteomics data have been deposited to the ProteomeXchange Consortium via the PRIDE (13) partner repository with the dataset identifier PXD056237 and 10.6019/PXD056237

### Protein functional annotation and graphic visualization

A sequence search against a *H. sapiens* database was performed to discard any peptides that exactly matched host peptide sequences (considering leucine/isoleucine to be equivalent). Using *H. sapiens* and *O. volvulus* databases, a similar search was completed for *Wolbachia* and virus results. The processed spectra and peptide count results are available for each condition and sample in tables S1 and S2, respectively. Functional annotations for all *O. volvulus* or *Wolbachia* proteins were assigned using results from InterProScan v5.59-91.0 to identify gene ontology classification and InterPro functional domains, and GhostKOALA v2.2 to assign KEGG (Kyoto Encyclopedia of Genes and Genomes) annotations (14–18). Additional protein annotation was performed using PANNZER and Sma3s (19, 20). Potentially secreted proteins were identified using SignalP v6.0, where any protein with a predicted signal peptide and fewer than 2 transmembrane domains was classified as secreted (21). Protein conservation data across nematodes, bacteria and host were quantified using BLAST (22). Functional enrichment was performed using the website http://webgestalt.org/ for KEGG and InterPro domains while GOSTATS v2.50 was used for GO (Gene ontology) “molecular function” child term enrichment. Enrichment was considered significant if the FDR-corrected p-values were ≤ 0.05, and at least 3 proteins were represented in the enriched group. Protein abundance as normalized spectral abundance factors (NSAFs) were used to reduce bias in quantification toward larger proteins (23, 24). NSAF data was plotted in Excel spreadsheets (Microsoft) to compare protein abundances between tissues and conditions, and to perform statistical analyses.

### Protein and antibody production

The sequence for OVOC8391 and its filarial orthologues were retrieved from WormBase Parasite (25). MegAlign Pro from DNA Star was used to align the sequences with OVOC8391 as the reference. Two portions of this large gene with low similarity to other filarial species were identified (S2 Fig.). Primers were designed for one of the low similarity regions and purchased from Integrated DNA Technology (IDT) (Coralville, IA, USA) (Table S3). PCR amplification was completed using blunt end PCR using Phusion High Fidelity DNA Polymerase (Thermo Fisher Scientific, Waltham, MA, US) with an annealing temperature of 50°C. To amplify, *O. volvulus* adult worm complementary DNA (cDNA) was used, and manufacturer instructions were followed for all kits used. Following PCR, the results were visualized on a 1% agarose gel. Once confirmed, the gene fragment was ligated into linearized pET100D plasmid (Invitrogen, Waltham, MA, USA) and transformed into the TOP10 *Escherichia coli* strain. Colony PCR was performed to check if the fragment was transformed correctly. Plasmid preparations were completed using Qiagen Plasmid Kit. Plasmid preparations were sent to be sequenced (Azenta, Burlington, MA, USA) with sequence analysis being completed using SeqManPro v17.4 (DNAStar).

After sequence confirmation, the plasmid was transformed into the BL21 *E. coli* strain for protein production. Cultures were grown at 37° C, 180 rpm in Luria Broth (Sigma, St. Louis, MO) with ampicillin at 50 ug/mL (GoldBio). Once they reached an OD600 over 0.6, the culture was induced with 1mM Isopropyl β-D-1-thiogalactopyranoside (IPTG) (GoldBio) and grown for an additional three hours. Cells were collected with centrifugation at 10,000 g for 15 minutes. Pellets were stored at −80°C until purification.

Pellets were suspended in a urea buffer with sodium phosphate at pH 8. The cell suspension rocked for 1 hour at room temperature. The suspension was sonicated to further break up the cells at 30 second intervals a total of three times. Benzonase (Sigma, St. Louis, MO) was added at 1μL for every 10mL of buffer. The suspension was centrifuged at 10000g for 15 minutes to collect the lysate. The lysate was poured over a column with cobalt His-affinity beads (Sigma, St. Louis, MO). The column was washed with the suspension buffer and protein was eluted in a urea and sodium phosphate buffer pH8 with 500mM imidazole. Protein was further purified using an Electro-Eluter (Bio-Rad, Hercules CA, USA). The collected elute was concentrated using an Amicon Ultra 3.5 kD MWCO cutoff filter (Millipore-Sigma, Burlington MA, USA). Protein purity was evaluated by SDS-PAGE electrophoresis followed by staining with SimplyBlue SafeStain (Invitrogen). Protein was quantified using a Qubit.

Three BALB/c mice were immunized with 20 µg of recombinant OVOC8391-Fragment 2 in complete Freund’s adjuvant. After 2 weeks, the mice were boosted with 20 µg of recombinant protein in incomplete Freund’s adjuvant. An additional boost was completed 2 weeks later with 20 µg of recombinant protein in incomplete Freund’s adjuvant. Sera was collected from the mice six days later.

### Immunohistochemical localization of OVOC8391-1 in adult *O. volvulus*

Embedded nodules with adult *O. volvulus* were sectioned at 5 µm and mounted on slides. After deparaffinization, the sections were blocked with 10 % bovine serum albumin (BSA) solution for 30 minutes at room temperature. The first polyclonal mouse antibody OVOC 8391-2 was applied in a 1:100 dilution in 0.1% BSA and left at room temperature for one hour or at 4° C overnight. Polyclonal anti-mouse IgG produced in rabbit was used at a 1:1000 dilution for 30 minutes at room temperature. Alkaline phosphatase-anti-alkaline phosphatase antibody (APAAP) produced in mouse was added to the section at a 1:40 dilution and left at room temperature for 30 minutes. Fast Red TR/Naphthol AS was used as chromogen and Mayer’s hematoxylin solution as counterstain. All reagents were provided by Millipore Sigma, St. Louis, MO, USA. Nodule section images were taken using Olympus DP70 microscope digital camera (Olympus, Tokyo, Japan).

## Results

### Morphological description of nodules with live female worms with neoplasms

Our study is based on an accurate histological analysis of worm tissue. Stained nodules slides retrieved were digitalized and evaluated by two independent microscopists. Both microscopists agreed on worm numbers, viability and fertility of worms, but detection of PN was only indirectly included in the analysis (11). For the current study, only nodules with a single live female worm with or without PN were used, to avoid potential contamination by other worms. *O. volvulus* females without neoplasm (HC) showed well-delimited organs like uterus, intestine, lateral cord and muscles and the presence of normal embryogenesis. This is defined as the presence of intact morula-, coiled-, pretzel- and stretched mf (Fig. 1A, red staining). PN female worms had cells invading the pseudocoelom with a high cytoplasm/nuclei ratio (Fig. 1B-E). Important parasite organ systems were still clearly recognizable, for example two uterine branches, the uterus wall and the intestine (Fig. 1B). One of the uterus branches can be present but is difficult to identify because of heavy colonization with the invasive PN cells (Fig. 1C-E). The high affinity for the H&E stain in the PN tissue is indicative of high, condensed DNA content, because of rapid cell division.

### Proteins detected in neoplasms and other worm tissues

In a first step, we determined the proteins present in the main tissue typers (PN, uterus, body wall, intestine and embryos) from cross-sections of PN or HC female *O. volvulus* worms. In total, 3,150 eukaryote proteins (2,618 proteins from *O. volvulus*; 443 proteins from *H. sapiens*; 29 proteins from cRap; and 60 decoy proteins) and 165 proteins from *Wolbachia* were identified (Tables S1 and S2). 2,325 *O. volvulus* derived proteins were detected in any sample but only 1,534 proteins were present in at least two replicates that also had two peptides in at least one tissue. Among these confidently identified proteins, 1,390 were present in the PN tissue with 2 peptides in 2 replicates (Fig. 2A). The total number of proteins present only in the PN tissue was 1,221, but if the proteins that were also detected in HC parasite tissues are disregarded, this number goes down to 594 PN unique proteins (Fig 2A). In the HC worms (Fig. 2B), 929 *O. volvulus*-derived proteins were detected, with 133 shared among all tissues. Embryo tissues (morula-, coiled-, pretzel- and stretched mf stages) of HC have with 574 unique proteins the most diverse proteome when compared with other HC tissues. Embryo tissue of HC contained 109 unique proteins that were not detected in PN worms. Because viruses can be linked to tumor development and an RNA virus that elicits an antibody response in humans was reported from *O. volvulus* (26) we searched our peptide results using a virus database. However, but no matches to viral proteins were found.

**Fig 2.**
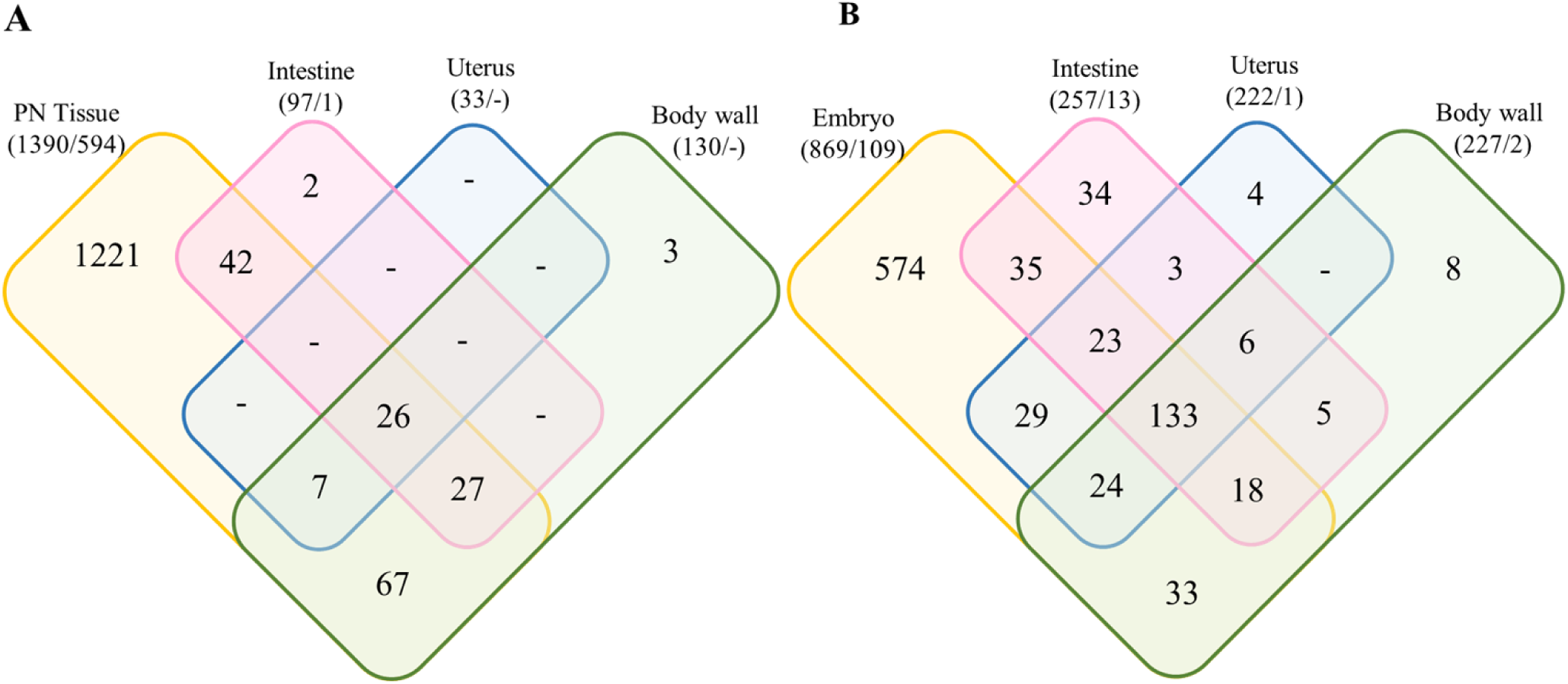
Venn diagram representing the number of *O. volvulus*-derived proteins for each tissue in the PN (A) and HC (B) worms. The overlap region between the circles shows proteins present in two or more stages. Numbers in parenthesis represents the total protein number found for the tissue/ the number of unique proteins not detected in the other sample type (ie, detected in PN but not in HC in (A) and in HC but not PN in (B)). PN: pleomorphic neoplasm, HC: healthy control.

### Proteins associated with adult worm neoplasms

In order to identify proteins or groups of proteins associated with PN, we compared the protein signature of PN with the proteins found in the intestine, body wall and uterus of the same worm and the corresponding tissues in HC worms (Fig 2). From the 1,390 *O. volvulus*-derived proteins found in PN tissue, 594 were not detected in any tissue of HC worms (Fig 2A and table S1). In addition, one interesting result was that several tissues looked healthy and were easily recognizable by shape in the PN worms, but fewer proteins were detected in each PN tissue than their counterpart in the HC worms (Fig 3). When compared, fewer proteins were identified in the intestine tissue from PN worms than in HC worms (97 vs 222, Fig 2) and they have only 66 proteins in common, representing 26.1% of all the proteins found in intestine tissues. Similar observations were made with the tissues of the uterus and body wall, where the number of overlapping proteins between PN and HC tissues was 12.7% and 45.5%, respectively (Fig 3).

**Figure 3.**
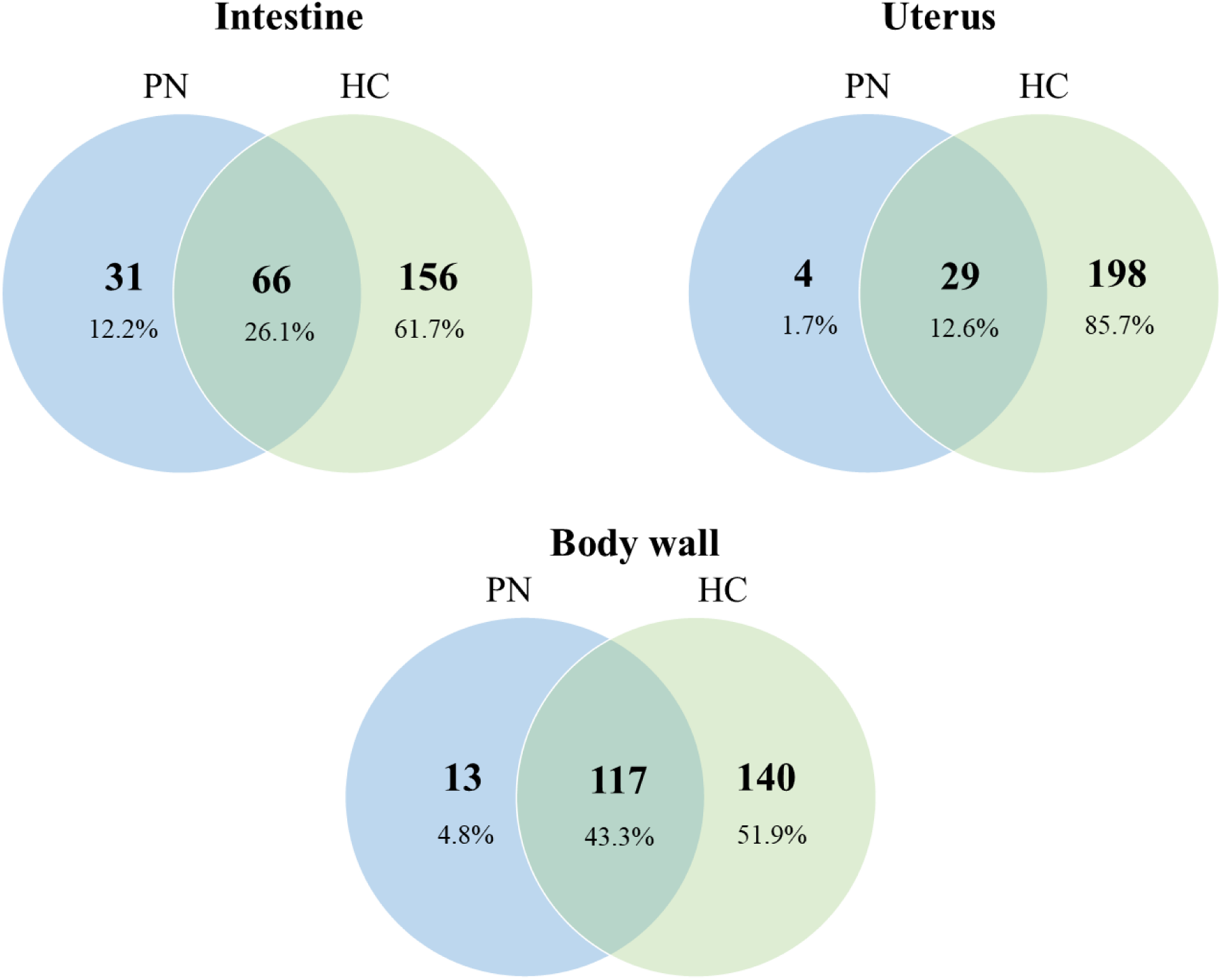
Comparison of protein from tissues from PN and HC O*. volvulus* worms. Venn diagrams represent the total number of proteins found for each tissue. Percentages are from the total protein PN: pleomorphic neoplasm, HC: healthy control.

We performed over-representation enrichment analysis in order to functionally characterize our different protein sets of interest from different tissues, using Gene Ontology (15) for functional term enrichment, KEGG (18) for pathway enrichment and InterPro (16) for functional domain enrichment. Table 3 shows the 10 most significantly enriched pathways identified for each database for the PN tissue. The complete list of enriched proteins is presented detailed in Table S4. Concerning the KEGG pathways, the most significant are for the ribosome, citrate cycle, glycolysis and proteasome pathways. Protein-protein enrichment analysis traced 10 InterPro domains; the categories with lowest FDR were nucleotide-binding alpha-beta plait domain superfamily, RNA-binding domain superfamily and RNA recognition motif domain. GO analysis identified 141 terms for cellular components with FDR-adjusted *P* values < 0.05, including the most significant terms “intracellular anatomical structure” and “cytoplasm”. Likewise, 75 different molecular functions GO terms were identified being RNA binding, structural molecule activity structural constituent of ribosome the domains with the most significant p-values. Similarly, 228 biological processes were significantly enriched and cellular amide metabolic, amide biosynthetic and peptide metabolic processes were the principal categories with lowest FDR-adjusted *P* value and more than 250 proteins included for each process.

**Table 3.** Significantly enriched gene ontology pathways, InterPro and KEGG domains. *O. volvulus*-derived proteins detected in the PN tissue, only proteins that were supported by at least 2 unique peptides in two biological replicates were used. The top 10 enriched terms for each functional category are shown.

| *GO domains – Biological Process* |  | *Total N°. O. volvulus detected* | *N°. PN proteins* | *FDR-adjusted P value* |
| --- | --- | --- | --- | --- |
| GO:0043603 | cellular amide metabolic process | 284 | 126 | 3.19E-20 |
| GO:0043604 | amide biosynthetic process | 261 | 119 | 3.19E-20 |
| GO:0006518 | peptide metabolic process | 271 | 119 | 1.11E-18 |
| GO:0043043 | peptide biosynthetic process | 255 | 114 | 1.11E-18 |
| GO:0006412 | translation | 251 | 112 | 2.51E-18 |
| GO:1901566 | organonitrogen compound biosynthetic process | 438 | 161 | 9.74E-17 |
| GO:0044281 | small molecule metabolic process | 314 | 117 | 4.61E-12 |
| GO:0034645 | cellular macromolecule biosynthetic process | 339 | 121 | 5.13E-11 |
| GO:0006082 | organic acid metabolic process | 158 | 66 | 9.12E-09 |
| GO:0055086 | nucleobase-containing small molecule metabolic process | 136 | 59 | 1.46E-08 |
| *GO domains - Molecular function* |  |  |  |  |
| GO:0003723 | RNA binding | 331 | 124 | 1.20E-13 |
| GO:0005198 | structural molecule activity | 229 | 94 | 4.56E-13 |
| GO:0003735 | structural constituent of ribosome | 131 | 62 | 1.07E-11 |
| GO:0051082 | unfolded protein binding | 28 | 21 | 1.34E-08 |
| GO:1901265 | nucleoside phosphate binding | 910 | 243 | 1.34E-08 |
| GO:0000166 | nucleotide binding | 910 | 243 | 1.34E-08 |
| GO:0036094 | small molecule binding | 989 | 258 | 3.61E-08 |
| GO:0004298 | threonine-type endopeptidase activity | 15 | 14 | 4.45E-08 |
| GO:0097159 | organic cyclic compound binding | 1921 | 446 | 1.40E-07 |
| GO:1901363 | heterocyclic compound binding | 1920 | 445 | 1.77E-07 |
| *GO domains - Cellular component* |  |  |  |  |
| GO:0005622 | intracellular anatomical structure | 1994 | 504 | 7.01E-71 |
| GO:0005737 | cytoplasm | 1051 | 296 | 3.20E-39 |
| GO:0043229 | intracellular organelle | 1709 | 406 | 3.64E-39 |
| GO:0043226 | organelle | 1738 | 408 | 4.45E-38 |
| GO:0043228 | non-membrane-bounded organelle | 514 | 179 | 6.07E-34 |
| GO:0043232 | intracellular non-membrane-bounded organelle | 514 | 179 | 6.07E-34 |
| GO:0032991 | protein-containing complex | 779 | 229 | 1.36E-31 |
| GO:0005840 | ribosome | 134 | 63 | 1.81E-18 |
| GO:0000502 | proteasome complex | 24 | 20 | 2.54E-12 |
| GO:0099080 | supramolecular complex | 126 | 52 | 2.54E-12 |
| *InterPro domains* |  |  |  |  |
| IPR012677 | Nucleotide-binding alpha-beta plait domain superfamily | 126 | 58 | 1.82E-11 |
| IPR035979 | RNA-binding domain superfamily | 126 | 57 | 3.98E-11 |
| IPR000504 | RNA recognition motif domain | 110 | 51 | 1.81E-10 |
| IPR001353 | Proteasome, subunit alpha/beta | 14 | 14 | 5.92E-09 |
| IPR012340 | Nucleic acid-binding, OB-fold | 54 | 28 | 1.13E-06 |
| IPR027417 | P-loop containing nucleoside triphosphate hydrolase | 405 | 113 | 1.38E-06 |
| IPR029055 | Nucleophile aminohydrolases, N-terminal | 20 | 15 | 3.74E-06 |
| IPR027410 | TCP-1-like chaperonin intermediate domain superfamily | 9 | 9 | 2.01E-05 |
| IPR027413 | GroEL-like equatorial domain superfamily | 9 | 9 | 2.01E-05 |
| IPR043129 | ATPase, nucleotide binding domain | 30 | 18 | 2.01E-05 |
| *KEGG domains* |  |  |  |  |
| 3011 | Ribosome | 145 | 62 | 1.23E-07 |
| 20 | Citrate cycle (TCA cycle) | 24 | 18 | 1.22E-06 |
| 10 | Glycolysis / Gluconeogenesis | 45 | 26 | 2.50E-06 |
| 3051 | Proteasome | 32 | 20 | 1.16E-05 |
| 3013 | RNA transport | 67 | 32 | 1.72E-05 |
| 3012 | Translation factors | 62 | 30 | 2.29E-05 |
| 3041 | Spliceosome | 173 | 58 | 8.93E-04 |
| 3110 | Chaperones and folding catalysts | 73 | 30 | 9.90E-04 |
| 970 | Aminoacyl-tRNA biosynthesis | 39 | 18 | 0.0052 |

Although the number of proteins found in the PN tissue was high, the abundance of these proteins was lower compared to other tissues where they were detected (Table S1). The most abundant protein was laminin (OVOC10067) which was also detected in all other tissue types. It was more abundant in the intestine, body wall and uterus for PN and HC worms but was less abundant in the embryos than in the NP tissue. Among the 10 most abundant proteins in the PN tissue, only two proteins did not share substantial sequence conservation with the host and are nematode specific: a pepsin inhibitor (OVOC9984) and a calcium binding protein (OVOC8391). Comparing the proteins results with the gene expression level in adult female worms (RPKM, reads per kilo base per million mapped reads), 73 proteins present in the PN tissue were not supported by expression detection in the previous RNA-seq analysis (10). From these, 24 belong to unique proteins in the PN tissue (Table S1). There are 487 proteins that are always more abundant in the PN tissue compared with all HC tissues studied here (Fig 4, yellow dots) and they are proteins that belong to pathways related to ribosomes, proteins processing and RNA transport (Table 3 and Table S1). For HC body wall, uterus and intestine 33, 88 and 38 of these proteins were found in these tissues, respectively (Fig 4A-C, yellow dots). When PN and HC embryo tissues are compared for these proteins, 296 proteins are more abundant in the PN tissue and 182 are only present in PN (Fig 4D, yellow dots). PN and HC embryo tissues share 420 proteins (Fig 4D, red dots) indicating that PN have more proteins in common with embryos than with any other tissue in the HC worms.

**Fig 4.**
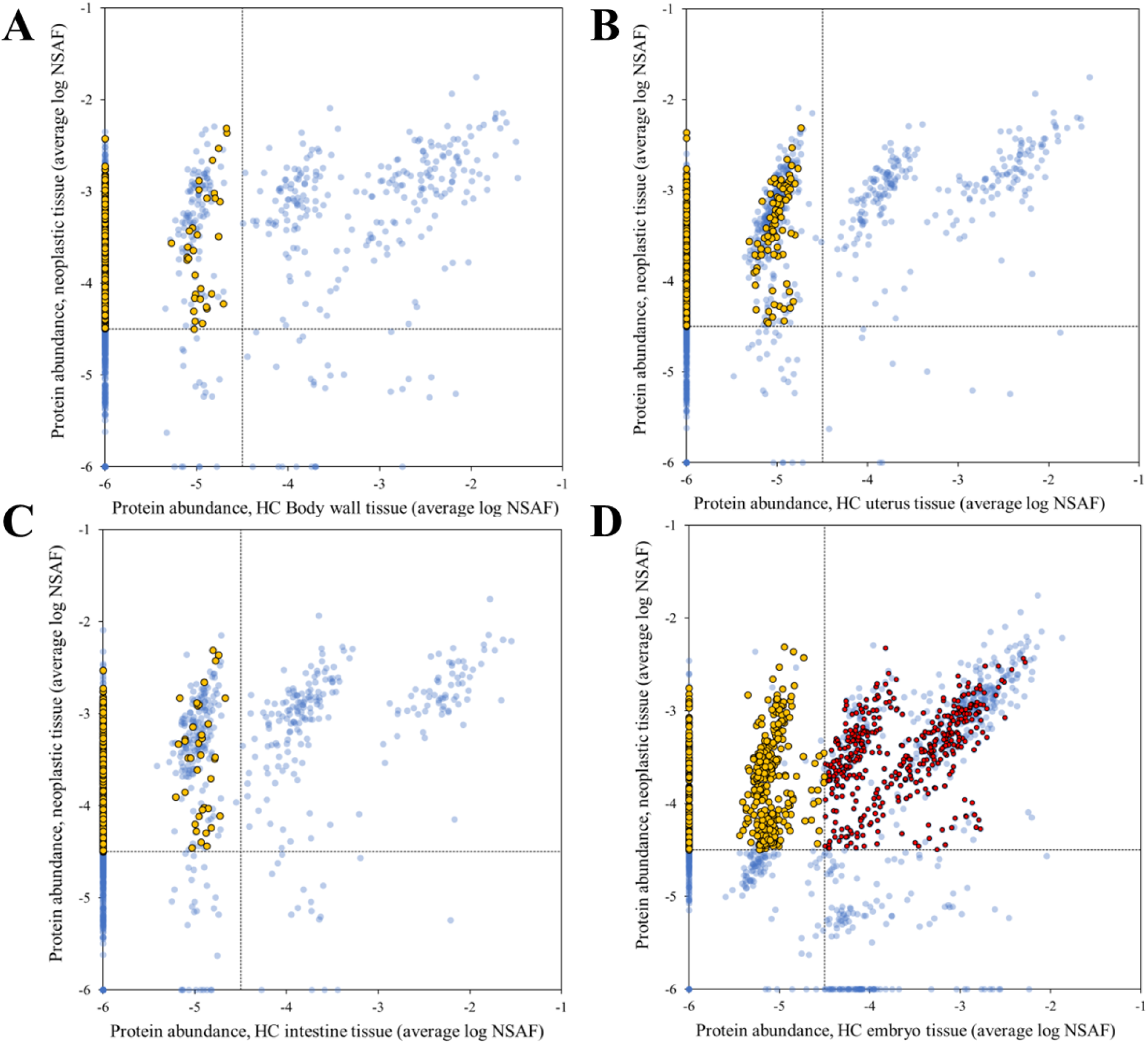
Protein abundance correlation between PN and HC worm tissues. The relative abundance of each detected protein in HC body wall tissue (A), HC uterus (B), HC intestine (C) or embryo tissue (D) proteins were plotted against the relative abundance in the PN tissue. Yellow dots represent proteins that are more abundant in the neoplasmic tissue compared to all HC tissues. Red dots are proteins that are only highly abundant in neoplasmic and HC embryo tissue when compared to all other HC tissues proteins. PN: pleomorphic neoplasm, HC: healthy control.

### Immunolocalization of the immunoglobulin-like protein *Ov*Dig-1

To confirm our proteomic results using an independent method, we decided to clone, express, and generate antibodies to the calcium binding protein OVOC8391 (*Ov*Dig-1). This large protein has a well-characterized ortholog in *Caenorhabditis elegans* (*Ce*Dig-1) in which this giant immunoglobulin-like protein (13,100 amino acids) is predicted to enable calcium ion binding activity (27). *Ce*Dig-1 is expressed in several structures, including glutamate-like receptors, body wall musculature, head mesodermal cells, male sex myoblasts, and non-striated muscles. In HC *O. volvulus* worms the protein was commonly detected by proteomics in all analyzed tissue and in PN worms it was found only in the body wall and NP tissues (Table S1). Immunolocalization confirmed these results, with *Ov*Dig-1 being highly expressed in intrauterine and pseudocoelomic PN, but distribution was linked to specific cell types (Fig 5). *Ov*Dig-1 was highly expressed in larger cells with high plasma content (Fig 5C-F). The protein was absent in the pseudocoelomic PN with a large number of small, dividing cells as indicated by a strong nuclear DNA stain (Fig 5E).

**Fig 5.**
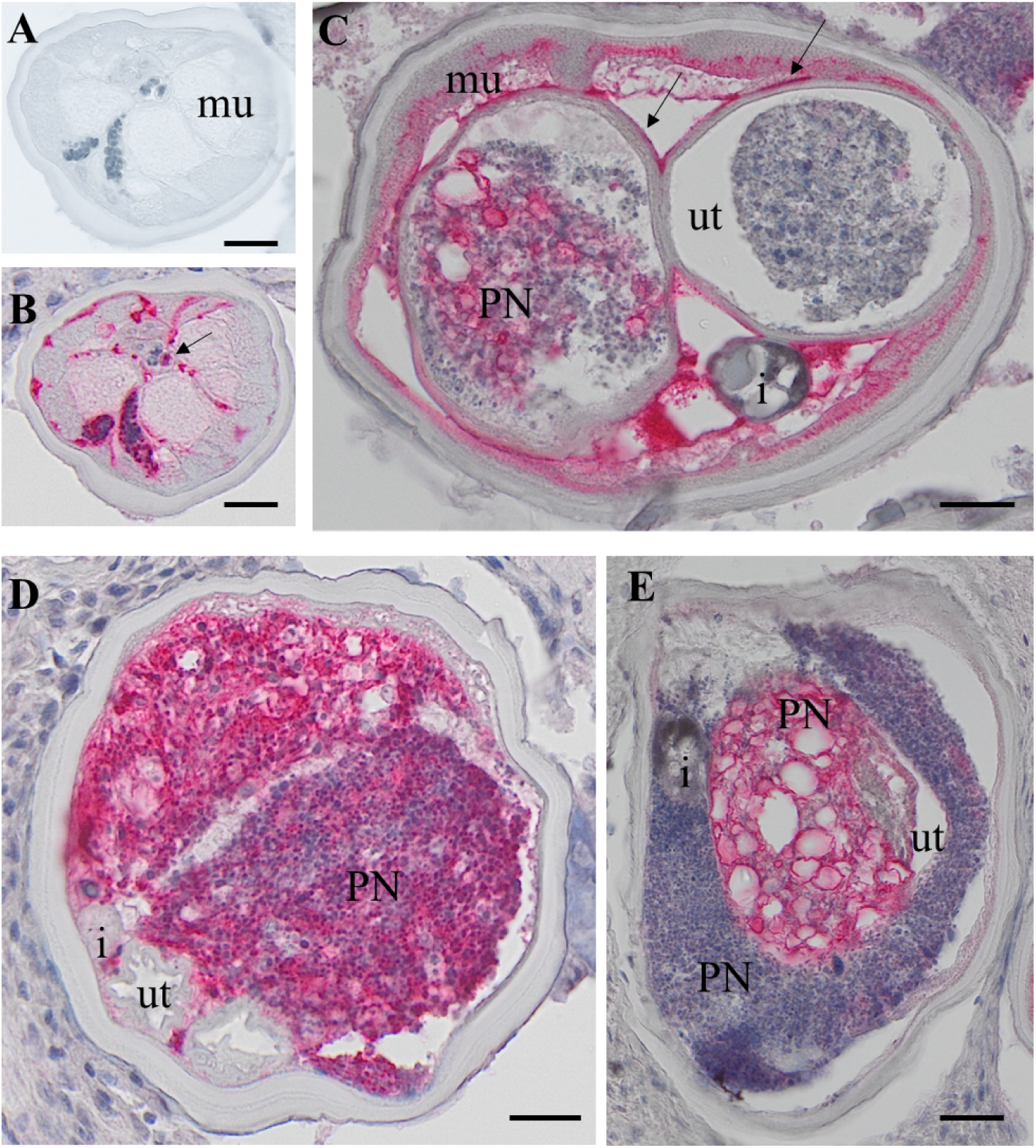
Immunhistolocalization of *Ov*Dig-1 (OVOC8391) in a PN female *O. volvulus*. A: Cross-section of a *O. volvulus* female with a few stretched Mf in the vagina, negative control using pre-immune serum. B: Same section as in A but stained (red) with an antiserum raised against *Ov*Dig-1. The protein is localized in the hypodermis, connective muscle tissue and in the neoplasm. Stretched Mf are also *Ov*Dig-1 positive (arrow). C: Cross-section of a female with PN cells in both uterus branches. The outer uterus wall, the connective tissue, muscle and PN cells are *Ov*Dig-1positive (red). D: PN in the pseudocoelom cavity with strong staining for OvDig-1. Both uterus branches are empty, constricted and not labeled. E: PN in the uterus strongly labeled for *Ov*Dig-1. PN in the pseudocoelomic cavity shows a more homogenous cell type with weak labeling. HC: healthy control, PN: pleomorphic neoplasm, i = intestine, mu= muscle, ut =uterus, scale bar= 50 μm.

In female *O. volvulus* with intact embryogenesis and without PN, *Ov*Dig-1 was expressed in primary oocytes, coiled and stretched microfilariae, connective tissue, outer uterus membrane and in the fibrillar zone of muscle cells (Fig 5). Red labelling for the protein was not observed in the rest of the muscle cells, the hypodermis or lateral chord.

### *Wolbachia*-derived proteins in healthy and pleomorphic neoplasm worms

*Wolbachia* proteins were curated the same way *O. volvulus* proteins and all peptides that may have originated from the mammalian host or the parasite were excluded from the final protein list (Table 4 and Table S2). Overall, relatively few *Wolbachia* proteins were detected in 2 of the 3 worms with PN. In the PN worms most *Wolbachia*-derived proteins were detected in the body wall (six proteins), followed by PN tissue (four proteins) and only one protein in the uterus. In the HC worms, eight *Wolbachia* proteins were detected in the body wall and seven in the embryos with an ATP-dependent chaperone (WP_025264048.1) found in both. Two *Wolbachia* proteins were found in the intestine but also in the embryo and body wall tissues. Body wall tissue from the NC and HC worms have all six proteins shared. Blast for all these hits match Rickettsiales bacteria and several arthropods (S2 Table).

**Table 4.**
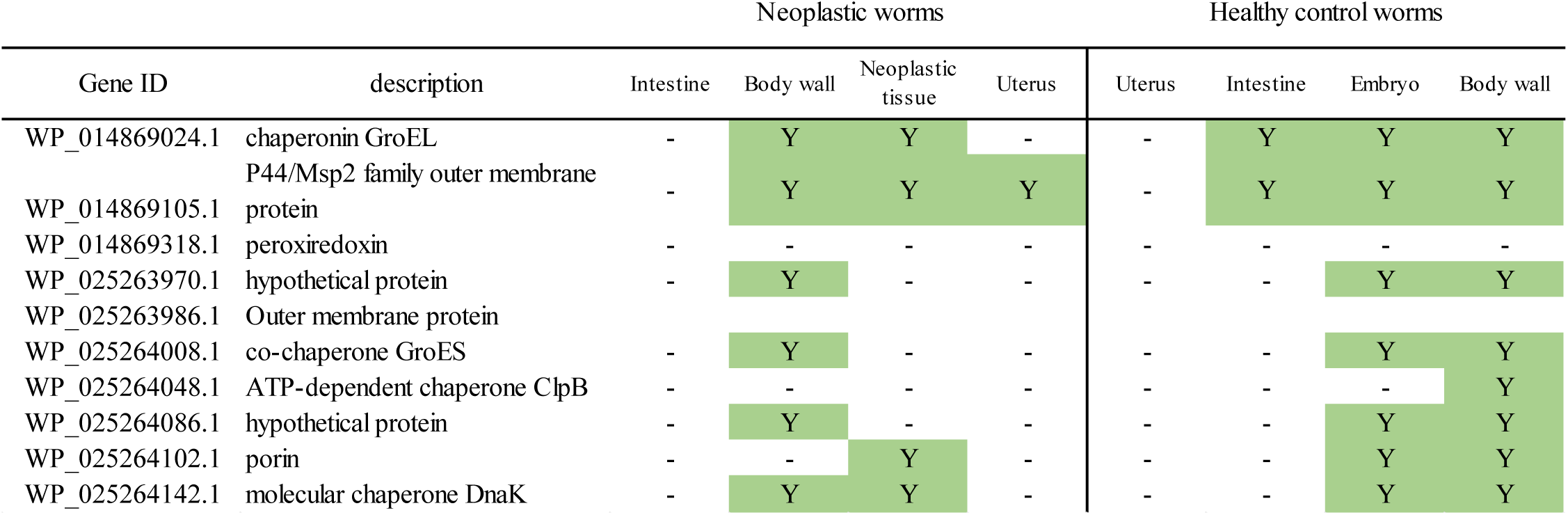
A summary of *Wolbachia* derived proteins detected in the *O. volvulus* samples. Presence of *Wolbachia-*derived proteins in the *O. volvulus* samples.

## Discussion

This study is the first proteomic analysis of *O. volvulus* worms with PN. PN in *O. volvulus* female worms were first reported by Duke *et al.* in 1990 and were more closely described in several subsequent studies (4, 5, 28). PN can occur in the pseudocoelomic cavity and uterus branches where they can be easily differentiated in histological sections from normal and degenerated embryos by their distinct morphology (Fig 1). PN were also found in adult male parasites, but they are less common than in females (4). Previous immunohistological studies of nodule sections indicated that PN are caused by filarial tissues and not by an invasion of human host cells. All eight filarial proteins examined by immunohistology previously were also detected in PN in our comprehensive LCM-LC/MS study (Table S1). In addition, we selected the nematode-specific calcium binding protein *Ov*Dig-1 (OVOC8391), that has a well-characterized ortholog in *C. elegans*, for expression, generation of antibodies and immunolocalization and were able to localize this protein in a specific cell type within the PN.

PN cells in *O. volvulus* adults appear first at the posterior end and spread from there to the anterior end. It was postulated that their origin could be in either ovarian or testis cells (4, 5). We studied only female worms, and ovaries are very short so that no cross-sections were found in PN worms that were suitable for proteomic analysis. However, we observed that 86% of the proteins found in HC embryo tissue (745 proteins) were also detected in PN tissue. Among these proteins, 296 were more abundant in PN and 420 proteins were abundant in both tissues (Fig 4D, yellow and red dots). The 296 more abundant proteins in PN are enriched for members of the protein synthesis pathways (ribosomes, proteins processing and RNA transport). Interestingly, these were the same pathways enriched for the proteins that are abundant in the PN and embryo tissue (Fig 4D, red dots and Table 3). Regulation of protein expression is a fundamental biological process during metazoan development and enriched regulatory proteins indicate rapid cell division. Regulatory proteins including transcription factors and the components of various signaling pathways are well-known for their roles in orchestrating organism development through transcriptional, translational or posttranslational control.

KEGG pathways, InterPro domains and GO terms enrichment for all the proteins present in the PN tissue indicated a strong enrichment of proteins associated with protein production specially with results related to ribosome, spliceosome, RNA transport and chaperones and folding catalysis (Table 3). Processes involved in protein degradation as proteasome and ubiquitin proteins are also enriched in this tissue. The PI3K-Akt signaling pathway is one of the most significantly enriched pathways, represented by 3 proteins in the PN, of the 5 proteins inferred from the *O. volvulus* genome, is one of the enriched proteins found in the neoplasm. This signaling pathway is considered mammals as a master regulator for cancer and is responsible for the regulation of cell growth, motility, survival, metabolism, and angiogenesis (29). Chromatin metabolism is frequently altered in cancer cells and facilitates cancer development and this affects mRNA transcription level, which involve histone-dependent chromatin access (30, 31). Alteration in histones were seen in the PN tissue, in terms of different protein detection; 12 histone proteins were found in the PN tissue while only six, four and one histone proteins were found in other tissues (HC embryo, intestine, body wall, respectively; Table S1). These findings showed that the neoplasmic is a deregulated tissue. All tissues but PN have lower protein numbers and abundance reduced when compared with the HC worms (Fig 2). This is expected as the worms are in the process of dying as the PN tissues are expanding through the other, normal tissues. Overall, we identified a substantial number of different proteins in PN, protein abundance detection was generally lower (Table S1).

To confirm our proteomic findings using an independent method, the localization of *Ov*Dig-1 in PN and HC worms was studied in more detail (Fig 5 and 6). The calcium binding protein *Ce*Dig-1 (OVOC8391 ortholog) is well-studied in *C. elegans* and is involved in the contacts between cellular surfaces and their environment (32). In *C. elegans* the expression in muscles surrounding the gonads was described previously. The distinct labeling for *Ov*Dig-1 of a plasma rich cell type in PN is a new finding and its function is not clear. Calcification of adult *O. volvulus* related to drug treatment or old age has been frequently observed previously (5, 28, 33–35). It can be hypothesized that *Ov*Dig-1 plays also a role in pathogenesis of worm calcification and PN are a risk factor for calcification.

**Fig 6.**
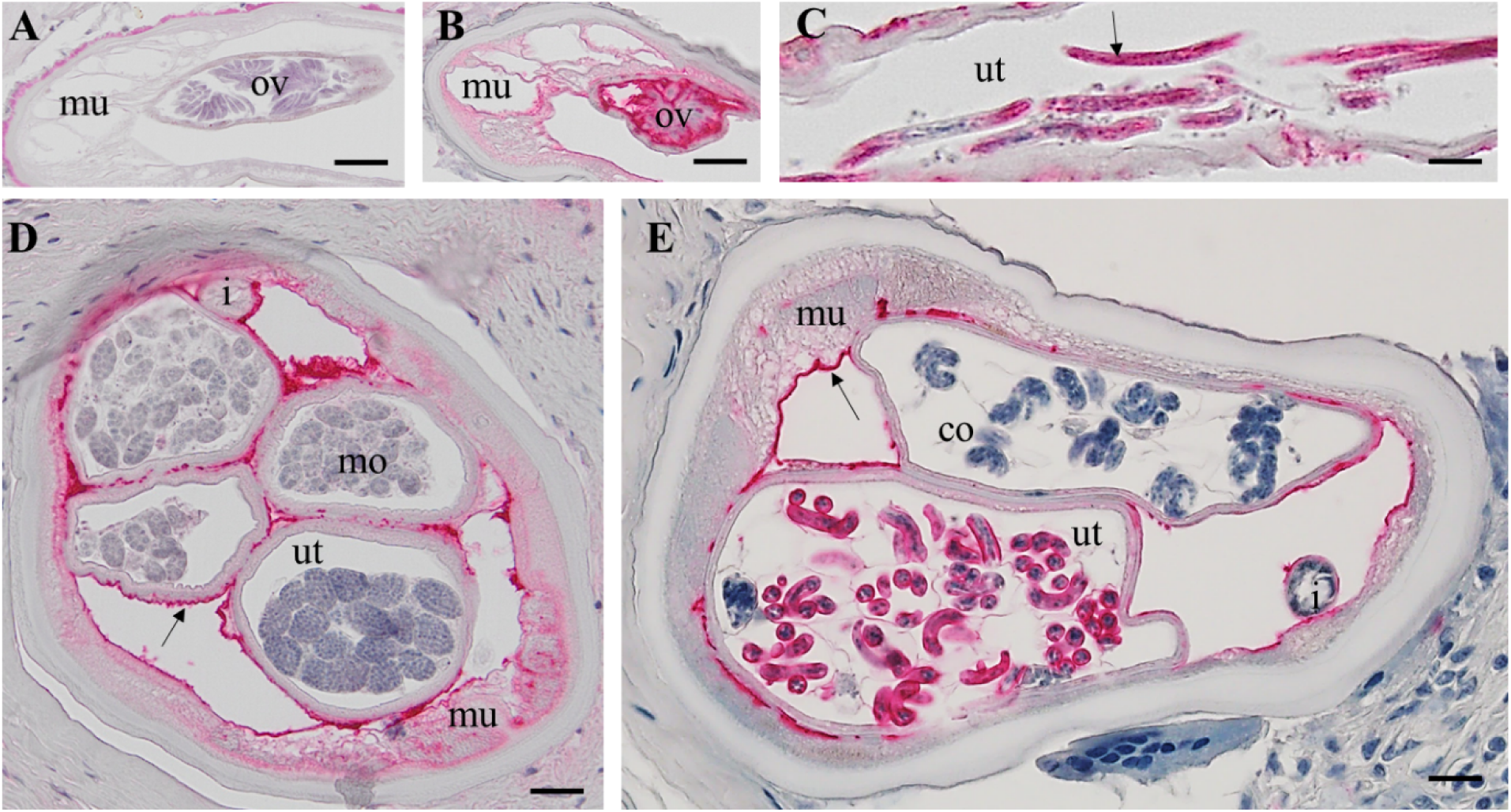
Immunohistolocalization of OvDig-1 (OVOC8391) in a healthy female *O. volvulus* worms. A: Non stained longitudinal section with primary oocytes in the ovary. B: Consecutive section as in A stained for *Ov*Dig-1 with positive primary oocytes (red). C: Longitudinal section of a uterus branch with stretched Mf that are strongly positive for *Ov*Dig-1. D: Cross-section of a female with different morulae stage embryos in the uterus. The outer uterus membrane is stained, some muscle cells and connective tissue. E: Cross-Section of a female with coiled (pretzel) stages in the uterus. More advanced coiled mf are OVOC8391 positive, while unmature mf are negative, like the morulae. co= coiled embryos i = intestine, lc = lateral chord, mo= morulae, mu= muscle, ov=oocytes, ut =uterus, scale bar= 20 μm

The combination of morphological and proteomic techniques can also lead to interesting and unexpected findings. A morphological difference between worms with PN and HC without is obvious, but a close comparison (see Fig. 5 and 6) shows no distinct morphological difference between structure such as the body wall or the intestine that are not directly affected by PN. However, our proteomic analysis shows a significant difference in protein composition in PN and HC worms also for those structures (Fig. 3 and table S1). For example, in the intestine of worms with PN only 97 proteins were detected with 31 unique intestinal proteins, while in the intestine of HC worms 222 proteins were detected with 156 unique intestinal proteins. This suggests that PN affect the entire worm and causes a systemic dysregulation of protein expression and is not confined to PN. This phenomenon is similar to early tumor metastasis in mammals, where primary-tumor-driven systemic processes that occur before metastasis have been found to dictate the site where subsequently disseminated cancer cells extravasate into other tissues (36).

*Wolbachia* is the most widespread genus of endosymbiotic bacteria in the animal kingdom, infecting a diverse range of arthropods and nematodes. Unlike many insects, filarial nematodes require *Wolbachia* for survival (37). In a female adult worm, *Wolbachia* can be observed in the hypodermis, lateral chord, oocytes and embryos. However, distribution in the hypodermis and the lateral chord is often uneven and *Wolbachia* density can vary. Brattig *et al.* observed *Wolbachia* frequently in worms with PN but not directly in the PN cells (4). In the present study we detected a small number of *Wolbachia* proteins directly in the PN cells (Table 4 and Figure S1). These proteins included the chaperonin GroEL, a surface protein, an outer membrane protein and a chaperone DnaK. The first 100 BLAST hits for these proteins shows a high similarity with the closely related *Rickettsiacae* and some arthropods species (Table S2). It is possible that these *Rickettsiales* sequences were mislabeled as arthropods/hosts sequences in the databases (38). The authors (us) discuss the possibility that these *Wolbachia* proteins were remnants of HC tissue cells or even off cuts from the adjacent tissues, but spectra count and peptide levels between neoplasmic and HC worms were too similar to consider that *Wolbachia* are in the sample by chance (Table S2). Overall, relatively few *Wolbachia* proteins were detected in the PN worms and the heathy control, but LCM was not targeting *Wolbachia* and the specific distribution of the endobacteria was not known.

## Conclusions

LCM combined with LC/MS is a powerful technology to assess the protein inventory of *O. volvulus* on a microscopic level. PN have a diverse proteome that is similar to the proteome of developing embryos in HC worms. The presence of PN is associated with a systemic dysregulation of protein expression also in organ systems that are not directly affected by neoplasms.

## Acknowledgements

The authors would like to express their gratitude to John Martin for his expert bioinformatics support during the proteomic analysis.

## Financial Disclosure Statement

This work was supported by grants INV-021433 (PUF) and OPP1190749 (GJW) from the Bill and Melinda Gates Foundation (https://www.gatesfoundation.org/) and by grant FBJH 5981 from the Foundation for Barnes-Jewish Hospital (https://www.foundationbarnesjewish.org/). The work at the Mass Spectrometry Technology Access Center at the McDonnell Genome Institute (MTAC@MGI) at Washington University School of Medicine, was supported in part by the Diabetes Research Center/NIH grant P30 DK020579, Institute of Clinical and Translational Sciences/NCATS CTSA award UL1 TR002345, and Siteman Cancer Center/NCI CCSG grant P30 CA091842.

The findings and conclusions in this paper are those of the authors and do not necessarily reflect positions or policies of the funding agencies. The funders had no role in the study design, data collection and analysis, decision to publish, or preparation of the manuscript.

## Supporting information

**Table S1.** Excel spreadsheet with the functional annotation, peptides, spectra count and NSAF values of the *Onchocerca volvulus* parasite-related-proteins and classification of the matched proteins by BLASTP searches against *Homo sapiens* and *O. volvulus* databases.

**Table S2.** Excel spreadsheet with the functional annotation, peptides, spectra count and NSAF values of the *Onchocerca volvulus* parasite-related-proteins and classification of the matched proteins by BLASTP searches against Wolbachia database.

**Table S3. Primers for OVOC8391 amplification**. Primers were designed for one of the low similarity with other parasites regions.

**Table S4. Complete list of enriched gene ontology pathways, InterPro and KEGG domains.** *O. volvulus*-derived proteins detected in the polymorphic neoplasm tissue, only proteins that were supported by at least 2 unique peptides in two biological replicates were used. Each tab represents one complete analysis.

**S1 Fig. Overview of H&E stained *O. volvulus* sections.** Worm sections which were used for LCM and which material was dissected using an image which had been manually color-coded to help keeping track which tissue should be dissected with the laser. A is a pleomorphic neoplasm worm and C is a healthy female. B and D are examples of images used at the LCM. Ut= uterus, b= bodywall, neo= neoplasm, m= male mo=morulae, Orange= neoplasm or embryos, green= gut; yellow= body wall, periwinkle= uterus wall.

**S2 Fig. Alignment for with filarial orthologues**. The sequence for OVOC8391 and its filarial orthologues were retrieved from WormBase Parasite. MegAlign Pro from DNA Star was used to align the sequences using OVOC8391 as reference.

## Notes

### Competing Interest Statement

The authors have declared no competing interest.

